# Neuroprotection following FLASH-RT may be mediated by sustained glutamate receptor AMPAR activation in CA3 neurons

**DOI:** 10.64898/2026.05.15.725423

**Authors:** Louis V. Kunz, Aymeric Almeida, Michèle Knol, Benoît Petit, Eniko A. Kramár, Marcelo A. Wood, Charles L. Limoli, Marie-Catherine Vozenin

## Abstract

To elucidate the early mechanisms underlying the long-term neuroprotective effect of FLASH-RT in the normal brain, spatial transcriptomics (Nanostring) were performed after whole-brain irradiation of C57BL/6J mice with either 1 or 3 fractions of 10 Gy at 5.6×10^6^ Gy/s (1 pulse-FLASH) or at conventional dose-rate 0.1 Gy/s. FLASH -RT induced a distinct transcriptomic signature in the CA3 and DG neurons, with upregulation of genes encoding glutamate receptors, involved in calcium signaling, long-term potentiation and mitochondrial OXPHOS. Early transcriptional upregulation of Gria gene translated into increased AMPAR protein levels at 48h in the DG and CA3 region and sustained higher AMPAR expression at 2 and 4 weeks post-FLASH. These findings support a durable activation of AMPAR. We propose a mechanism to explain FLASH-induced neuroprotection initiated by early calcium influx and subsequent sustained expression of glutamate receptor AMPAR in neurons and/or neural progenitors of the CA3, potentially contributing to long-term cognitive sparing.

**Graphical abstract:** 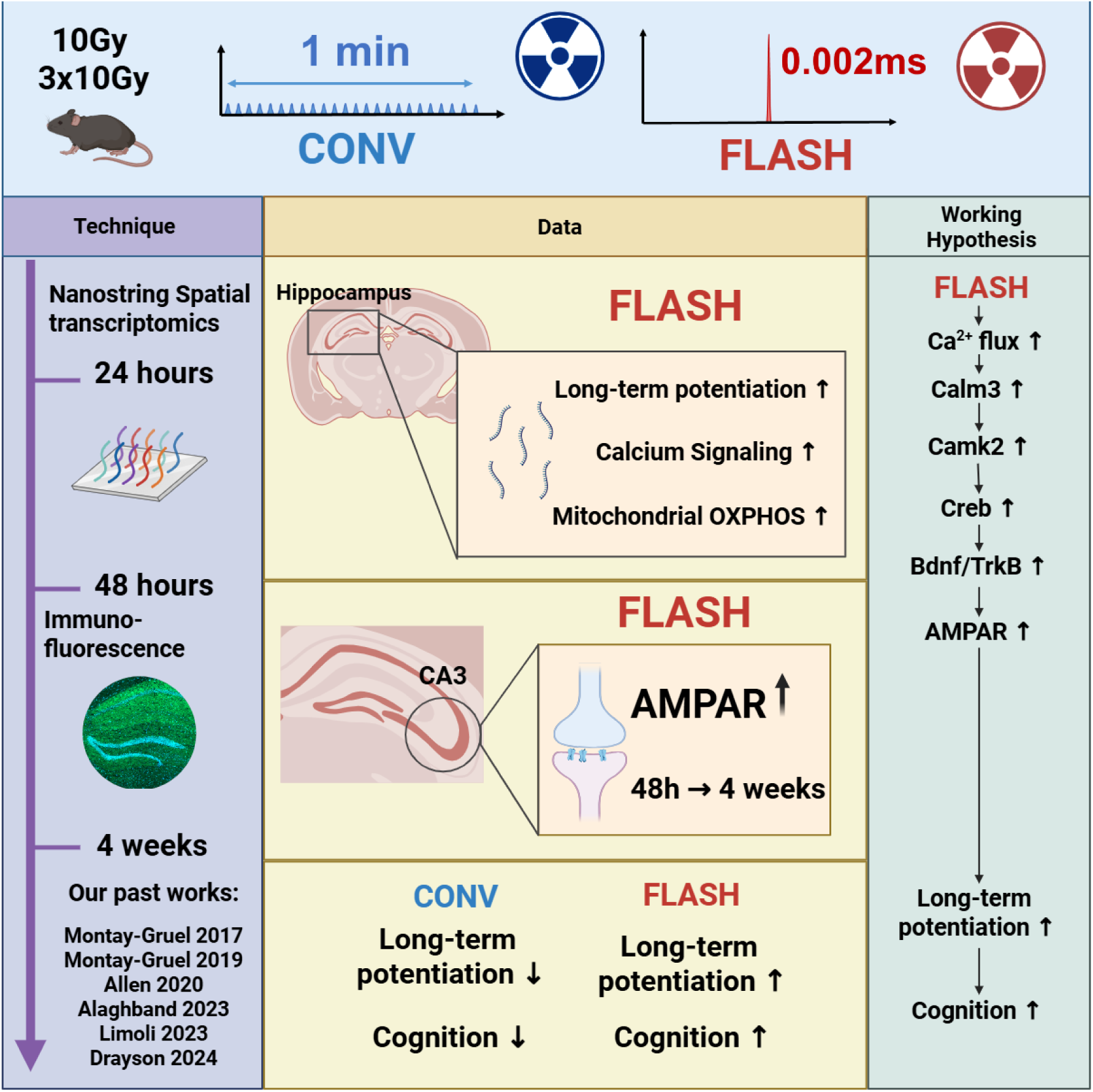

**Highlights:**

- FLASH-RT induces a stronger transcriptional response in the hippocampus than the cortex.
- FLASH-RT induces calcium signaling, LTP and mitochondrial OXPHOS genes.
- Early AMPAR upregulation leads to sustained protein expression.
- FLASH-RT induces a AMPAR-dependent signaling program in CA3 neurons.

## Introduction

With the dual benefit of mitigating radiation-induced normal tissue toxicity while preserving effective anti-tumor responses, FLASH radiotherapy (FLASH) represents a possible groundbreaking advancement in radiation oncology. This unique biological phenomenon, referred to as the “FLASH effect,” has been extensively investigated in the brain^1–8^ and reproduced across various tissues and species, using different radiation sources and delivery regimens ^9^.

The neurotoxic effects of standard cranial radiotherapy are well characterized. They involve cognitive decline, memory impairment, fatigue, white matter damages, neuroinflammation, vascular injury, and in severe cases radiation necrosis. Largely due to neuronal loss, demyelination, and disruption of neurogenesis in sensitive brain regions, these damages are irreversible, limit the dose that can be delivered to cure brain tumors and alter dramatically patients quality of life ^10,11^. While FLASH irradiation is not yet ready for clinical administration, preclinical studies done in rodents show that all the classical side effect of brain irradiation can be minimized by FLASH ^1–4,7,8^. For instance, FLASH irradiation does not alter dendrites, nor dendritic spines in neuron. Perforated synapses, that are believed to represent the structural correlates of learning and memory ^12,13^, are preserved after FLASH exposure ^3^, suggesting FLASH might spare structural proteins essential for cognitive preservation. Consistently, electrophysiological assessments of long-term potentiation (LTP) in the hippocampus and medial prefrontal cortex revealed that this metric of synaptic plasticity was preserved by FLASH but was permanently altered by conventional-dose rate radiotherapy (CONV) at late time-points post-irradiation ^5,6^. These findings show that at isodose, FLASH and CONV generate distinct effects on neuronal structure and function that might be transcriptionally regulated at early time point.

Considering the foregoing, we sought to investigate this question using early spatial transcriptomics and contrasted the imprint obtained after FLASH (5.6 MGy/s), CONV (0.1 Gy/s) and non-irradiated samples. Analysis focused on neurons over various brain regions including the hippocampal (HIP) subregions cornu ammonis region 1 (CA1, post-synaptic), region 3 (CA3, pre-synaptic), dentate gyrus (DG), stratum oriens (SO) as well as the cortex (CTX). Two treatment regimens that have been shown to preserve cognition in mice after FLASH compared to CONV were used: whole brain irradiation either in a single dose of 10 Gy or in a hypofractionated regimen composed of 3 fractions of 10 Gy, the latest being closest to clinical regimens. Brains were collected 24 hours after completion of irradiation to determine early transcriptional imprint.

## Results

### Transcription is not turned off by FLASH in the hippocampus

Given the intimate involvement of the cortex and hippocampus in learning and memory processing, early transcriptional modifications were investigated in these two brain regions by exposing the head of mice to a single or three fractions of 10 Gy delivered FLASH or CONV (Figure 1A). Unsupervised analysis using all quantifiable genes did not show any clear clustering (Figure supp 1D, 1E), but gene set enrichment analysis (GSEA) was more informative when performed on all the differentially expressed gene (DGE) sets. Venn diagrams were generated to visualize pathway overlaps between the various conditions with upregulated (UP) in red and downregulated (DOWN) in blue. (Figure 1B, 1C).

**Fig. 1.**
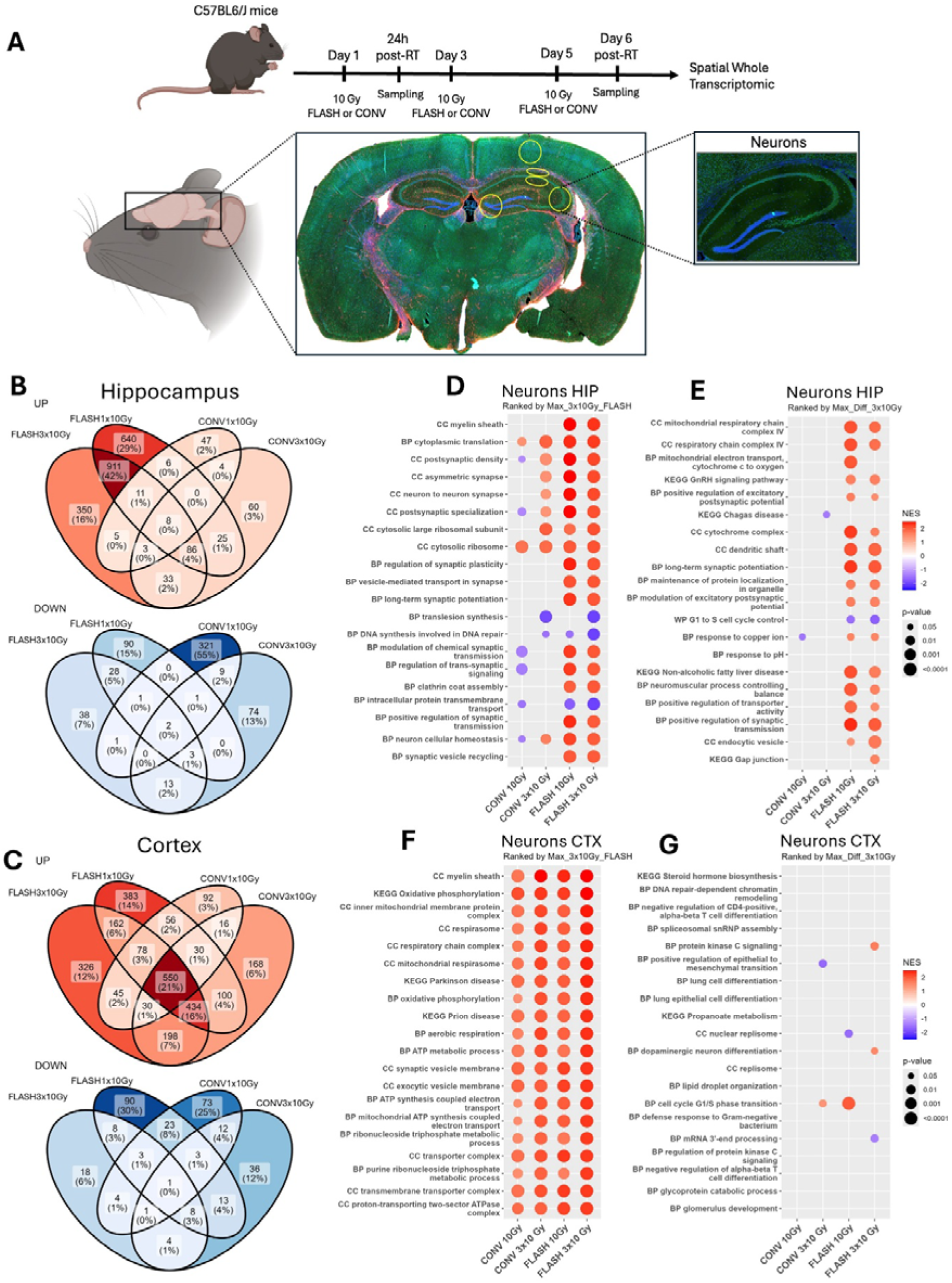
Cortical and hippocampal neurons show a different pattern of response to CONV, but not to FLASH. (A) Experimental design to investigate FLASH-specific genomic signature in mouse brain. (B) Venn Diagrams of Up (NES > 0, pvalue < 0.05) and Down (NES < 0, p-value < 0.05) regulated pathways in the Hippocampus (C) Venn Diagrams of Up (NES > 0, p-value < 0.05) and Down (NES < 0, p-value < 0.05) regulated pathways in the Cortex (D) Dotplot of pathways with all conditions compared to controls (CONV 10 Gy, CONV 3×10 Gy, FLASH 10 Gy, FLASH 3×10 Gy) in the Hippocampus, Normalized enrichment score (NES) as the colour and p-value as the dot-size, ranked according to the maximum average NES of FLASH 3×10 Gy (E) or the maximum absolute difference in NES between CONV 3×10 Gy and FLASH 3×10 Gy. (F) Dotplot of pathways with all conditions compared to controls (CONV 10 Gy, CONV 3×10 Gy, FLASH 10 Gy, FLASH 3×10 Gy) in the Cortex, Normalized enrichment score (NES) as the colour and p-value as the dot-size, ranked according to the maximum average NES of FLASH 3×10 Gy (G) or the maximum absolute difference in NES between CONV 3×10 Gy and FLASH 3×10 Gy. See also supp figure 1.

<u>In hippocampal neurons</u>, FLASH irradiation (both 1×10 Gy and 3×10 Gy) induced a markedly higher number of overexpressed pathways compared to CONV. Notably, the two FLASH regimens accounted for more than 50% (1551 and 1261 pathways, respectively) of the upregulated pathways, while CONV conditions induced relatively few unique pathways (2–4 %). Conversely, the downregulated pathways were predominantly associated with CONV 1×10 Gy (55% or 321 pathways), whereas FLASH (both 1×10 Gy and 3×10 Gy) contributed comparatively fewer unique downregulated pathways (5-15 %) (Figure 1B). Next, pathways were ranked either by maximum enrichment score in FLASH 3×10 Gy (Figure 1D) or by the largest difference in score between CONV and FLASH 3×10 Gy (Figure 1E). Strong induction of pathways associated with myelin sheath, synaptic plasticity, long-term synaptic potentiation, mitochondrial respiratory chain, respirasome, and the synaptic vesicle cycle were <u>uniquely</u> found in FLASH samples.

<u>In the cortex</u>, a substantial number of pathways were <u>shared</u> between FLASH and CONV condition (21%, 550 pathways) (Figure 1C). FLASH conditions still resulted in upregulation of several pathways, representing 14% (FLASH 1×10 Gy, 383 pathways) and 12% (FLASH 3×10 Gy, 326 pathways) of the total pathways. Pathways associated with mitochondrial respirasome, electron transport chain, oxidative phosphorylation, and synaptic vesicle components were upregulated both in FLASH and CONV samples (Figure 1F, 1G).

In summary, in the hippocampus a differential transcriptional pattern is found, with FLASH inducing a stronger transcriptional activation, while CONV is associated with more prominent gene downregulation relative to control. On the other hand, in the cortex, a more general radiation-induced imprint is found. This result reveals different transcriptional regulations in the hippocampus and cortex after FLASH exposure.

### FLASH induces a transcriptional program prone to neurotransmission in the hippocampus

FLASH elicits an hippocampal transcriptional program that closely resembles the radiation-induced program activated in the cortex independently of dose and dose rate. It involves up-regulation of pathways associated with myelin sheath, synaptic plasticity, long-term synaptic potentiation, mitochondrial respiratory chain, respirasome, and the synaptic vesicle cycle. In addition, FLASH induces a highly conserved transcriptomic signature that is largely dose-independent, in contrast to CONV irradiation where gene expression changes appear more dose-dependent (Figure 2A and 2B). This suggests that the major transcriptional response to FLASH is established after the first fraction of 10 Gy, remaining stable over the course of fractionation (Supplementary Figure 2A, 2B). In contrast, CONV irradiation halts transcription and the few induced pathways are related to stress and show a dose-dependent response, such as cell death after the first fraction of 10 Gy and translation, post-replication repair and oxidative stress response after the third fraction of 10 Gy, indicating clear cumulative damages (Supplementary Figure 2C, 2D).

**Fig. 2.**
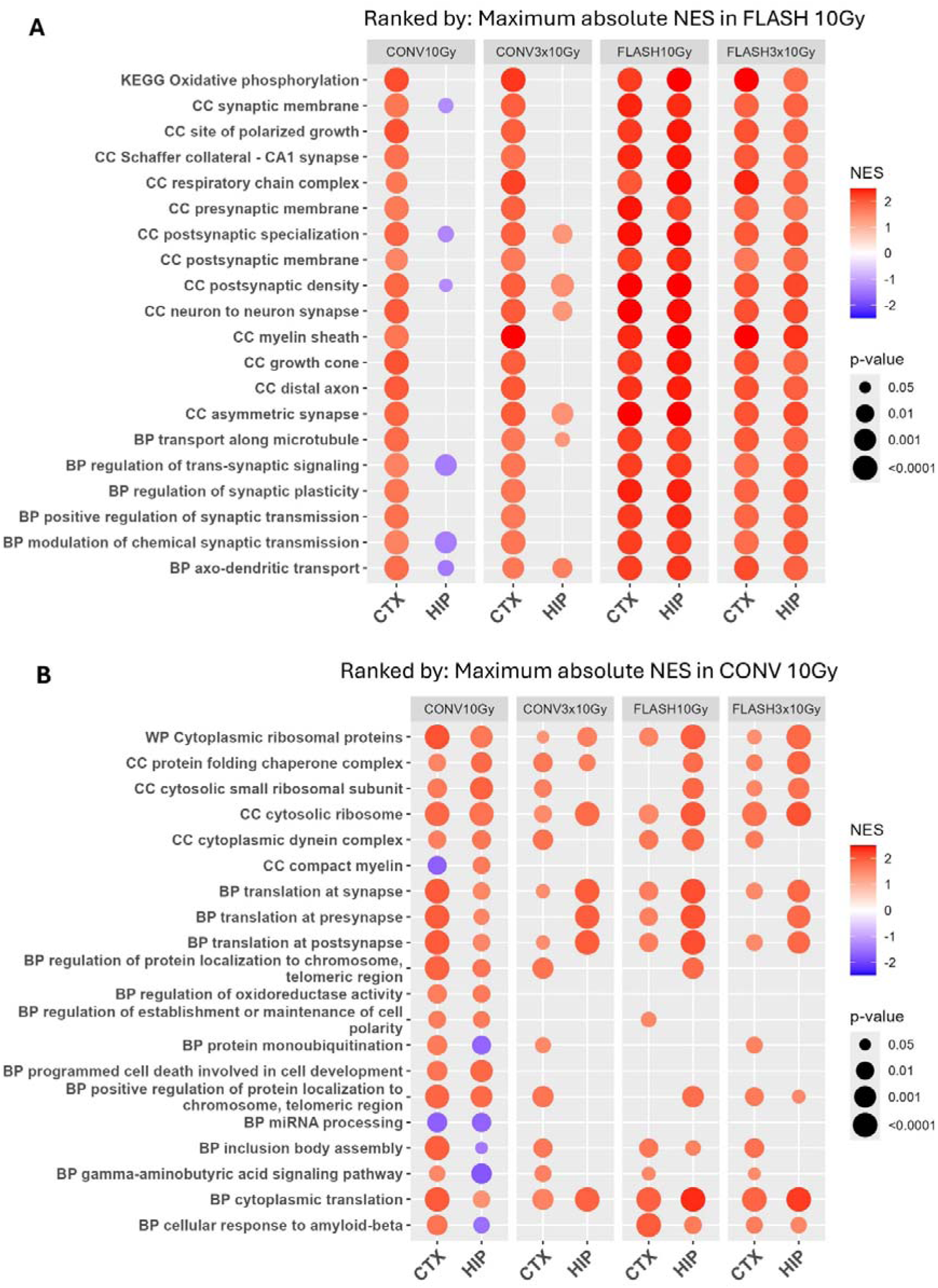
In the hippocampus, FLASH but not CONV induces a transcriptional program reminiscent of the cortex, independently of the total dose. (A) Top 20 pathways (highest absolute average NES across regions) after FLASH 10 Gy (B) and after CONV 10 Gy. Normalized enrichment score (NES) as the color and p-value as the dot-size. See also supp figure 2.

### Genes associated with synaptic function are overexpressed in FLASH-exposed CA3 and DG

UMAP algorithm revealed distinct clustering by region (Figure 3A) and showed that various neuronal populations display specific transcriptional profiles indicative of region-specific morphology, function and stress response including radiation. Therefore, transcriptomic changes were further analyzed separately across the CA1, CA3, DG, and SO subregions (Figure 3, Figure supp 3).

**Fig. 3.**
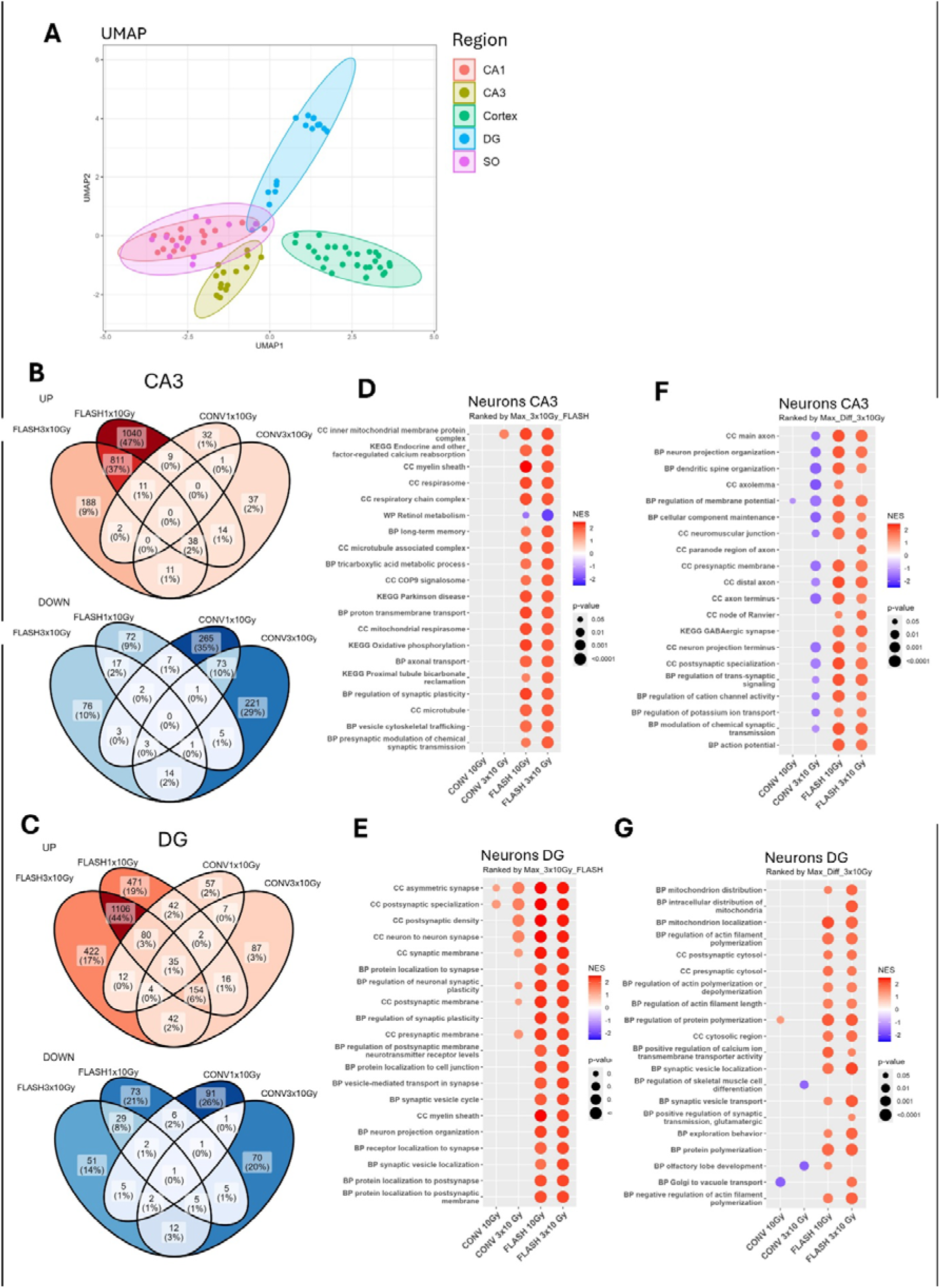
Hippocampal DG and CA3 regions are essential in the FLASH sparing effect. (A) UMAP and tSNE of neuronal transcriptomic signature with subregions colored and ellipses generated using *stat_ellipse*. (B) Venn Diagrams of Up (NES > 0, p-value < 0.05) and Down (NES < 0, p-value < 0.05) regulated pathways in the CA3. (C) Output of GSEA on CA3 signature, pathways are ranked according to the maximum average NES of FLASH 3×10 Gy (D) or the maximum absolute difference in NES between CONV 3×10 Gy and FLASH 3×10 Gy (E) Venn Diagrams of Up (NES > 0, p-value < 0.05) and Down (NES < 0, p-value < 0.05) regulated pathways in the DG. (F) Output of GSEA on CA3 signature, pathways are ranked according to the maximum average NES of FLASH 3×10 Gy (G) or the maximum absolute difference in NES between CONV 3×10 Gy and FLASH 3×10 Gy. See also supp figure 3.

Venn diagrams and GSEA showed that a substantial proportion of upregulated pathways were shared between FLASH 1×10 Gy and 3×10 Gy regimen in the CA3 and DG subregions, accounting for 37% and 44% of the upregulated pathways, respectively (Figure 3B, 3C). In contrast, fewer shared pathways were observed in the CA1 (18%) and SO (7%) subregions between the two FLASH conditions (Figure supp 3C, 3F).

Pathways were ranked by the highest enrichment score in FLASH 3×10 Gy (Figure 3D, 3E), FLASH 1×10 Gy (Figure supp 3A, 3B) or by the greatest differences in enrichment score between FLASH and CONV 3×10 Gy (Figure 3F, 3G) which highlighted that FLASH uniquely led to marked upregulation of pathways involved in myelin sheath formation, oxidative phosphorylation, neuronal organization, membrane potential regulation, synaptic transmission, vesicle cycling, actin polymerization, and calcium regulation when compared to unirradiated controls. These effects were <u>independent of dose</u>. Some pathways were also enriched in the CA1 and SO subregions, but a more pronounced dose dependency was observed in these regions (Figure supp 3).

To gain further molecular insight into the protective impact of FLASH on KEGG LTP pathway and KEGG synaptic vesicle cycle, illustrations of the pathways were generated (Figure 4, Figure supp 4A, 4B). Genes coding for AMPAR (*Gria2*), NMDAR (*Grin1*), and proteins essential for synaptic growth (*Nrxn1*, *Pfn2*, *App*) were found to be upregulated after FLASH. Genes involved in synaptic vesicle exocytosis such as *Snap25*, *Vamp*, *Syntaxin*, *Syt* and endocytosis such as *Dynamin*, *Clathrin*, *Ap2* were significantly upregulated in FLASH samples compared to CONV.

**Fig. 4.**
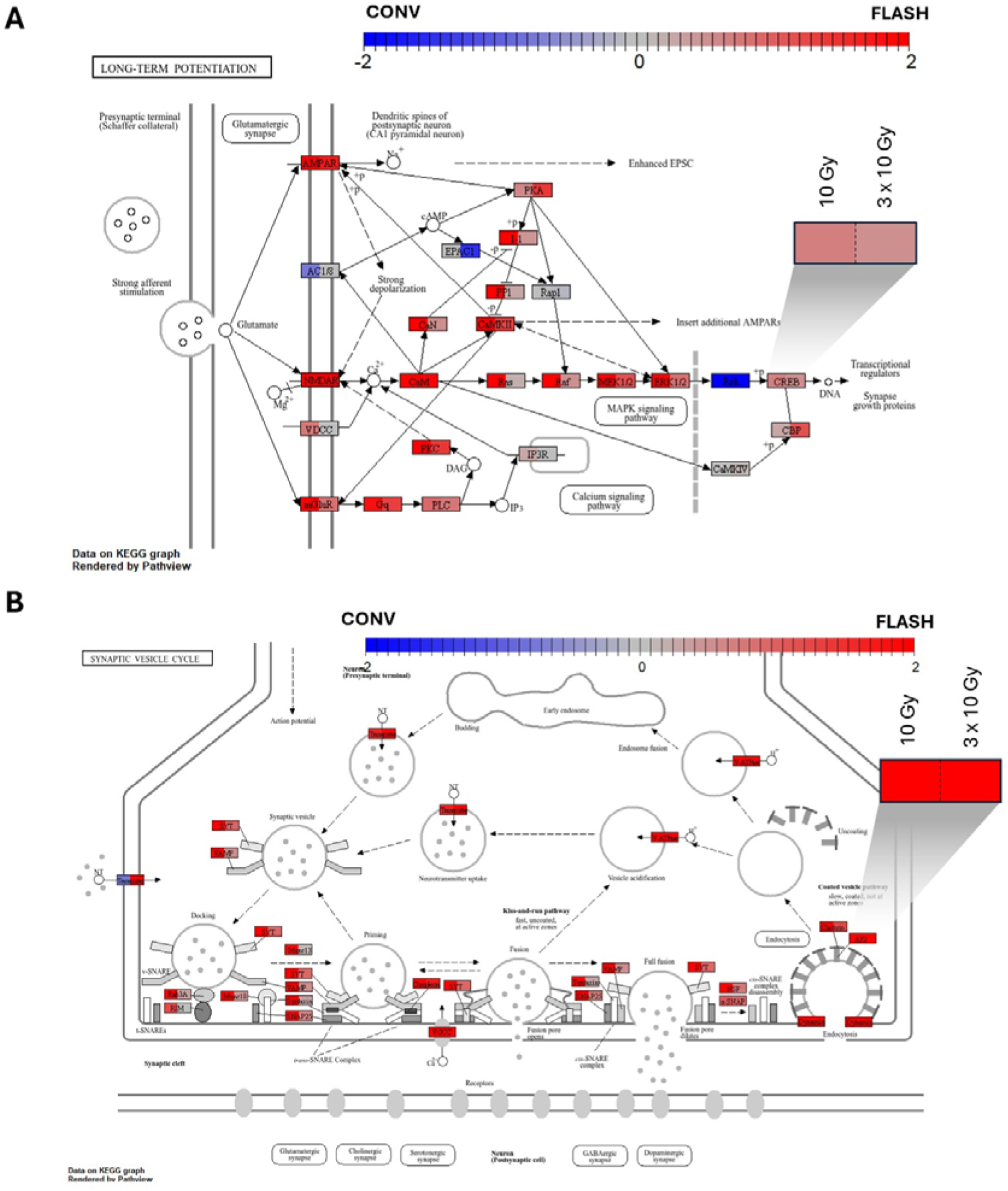
Genes associated with long-term potentiation and synaptic vesicle pathways are upregulated after FLASH but not CONV. (A) Pathways diagrams made using *Pathview* for KEGG long-term potentiation (B) and synaptic vesicle cycle based on CA3 imprint. Color is an arbitrary scale of enrichment from the highest expression in CONV (-2) to the highest expression in FLASH (+2). Each element of the pathway is split across center for CONV/FLASH comparison at 10 Gy (left) and at 3×10 Gy (right). See also supp figure 4.

### Preservation of glutamate receptors and calcium signaling might mediate the FLASH effect

Based on these findings, the rest of the analysis was focused on long-term potentiation (LTP) and synaptic vesicle cycle pathways, in the neurons of the CA3 and DG (Figure 5, Figure supp 4C, 4D). Our objective was to identify potential FLASH-specific genes associated with the sparing of LTP and possibly cognition preservation.

**Fig. 5.**
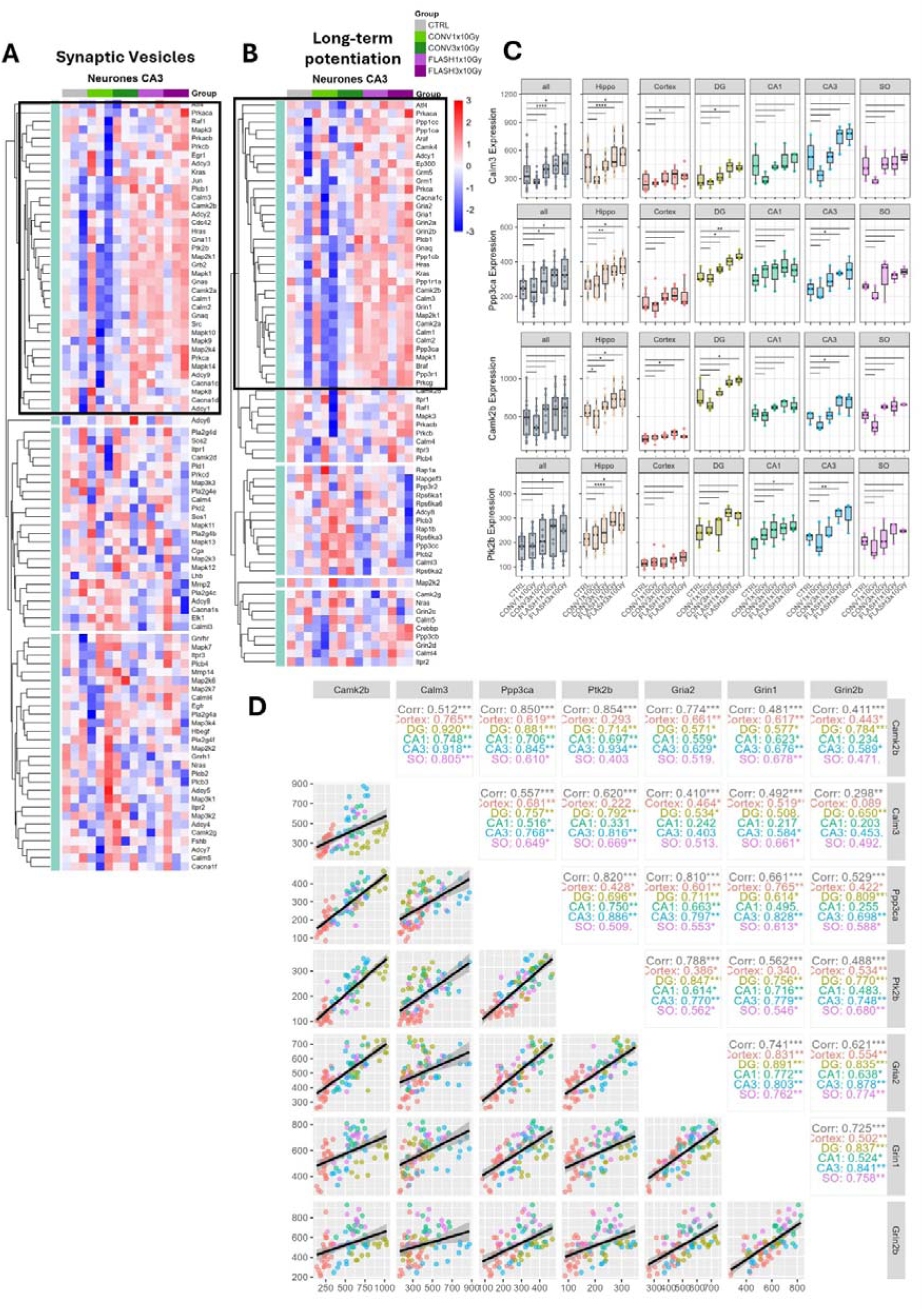
Expression of specific intercorrelated genes drives the long-term potentiation and synaptic vesicle cycle pathways. (A) Gene regulation heatmap for neurons of CA3, with black squares drawn arbitrarily for genes in pathways from KEGG database *synaptic vesicles* (B) and *long-term potentiation*. (C) Box plots of selected genes found in both pathways overall (all) or specifically in the hippocampus (Hippo), CA1, CA3, Cortex, DG or SO. One, two, three and four stars respectively representing p-values smaller than 0.05, 0.01, 0.001, 0.0001 as computed with the LMM model. (D) Correlation plot of selected genes showing a linear model fitted to all points using the lm function of R, along with correlation scores and p-values, computed with the same function. Selected genes are the same as above and subunits of AMPAR (*Gria2*) and NMDAR (*Grin1*, *Grin2b*). See also supp figure 4.

When the expression of genes involved in long-term potentiation and synaptic vesicle signaling was plotted as a heatmap and clustered by similarity, distinct clusters (black squares) emerged, showing genes that were indeed differentially upregulated following FLASH compared to CONV across both pathways and subregions (Figure 5A, 5B and Figure supp 4C, 4D). In the LTP pathway, a key cluster of upregulated genes was identified in each subregion after FLASH, regardless of the dose. Among the genes found in this cluster, *Calm3*, and *Ppp3ca* genes were consistently upregulated. Similar patterns were observed in the synaptic vesicle cycle pathway, with *Camk2b* and *Ptk2b* being significantly upregulated after FLASH. Intriguingly, *Calm3*, *Ppp3ca*, *Camk2b*, and *Ptk2b* exhibited strong inter-correlation in the hippocampus, possibly indicating a cooperative role in the protective effects of FLASH on LTP and cognitive function (Figure 5D). Quantification of Camk2 and phosphorylated Camk2 by immunofluorescent staining on mice hippocampi sampled 48h, 2 weeks and 4 weeks post irradiation showed that the ratio of pCamk2/Camk2 was similar in all groups (Figure supp 5). These calcium-signaling-related genes also showed good correlation with *Gria2*, *Grin1* and *Grin2b* genes coding for subunits of AMPAR and NMDAR. (Figure 5D). Immunofluorescence staining performed on brains sampled 48 hours post treatment showed a trend for AMPAR overexpression after FLASH in the CA3 and DG. This trend is maintained at 2 and 4 weeks in the CA3 only. A linear mixed model (LMM) considering all three timepoints confirmed that AMPAR is overexpressed in the FLASH irradiated animals compared to the conventionally irradiated animals (p-value<0.05) (Figure 6). Meanwhile NMDAR showed similar levels across conditions for the three timepoints investigated (Figure supp 6).

**Fig. 6.**
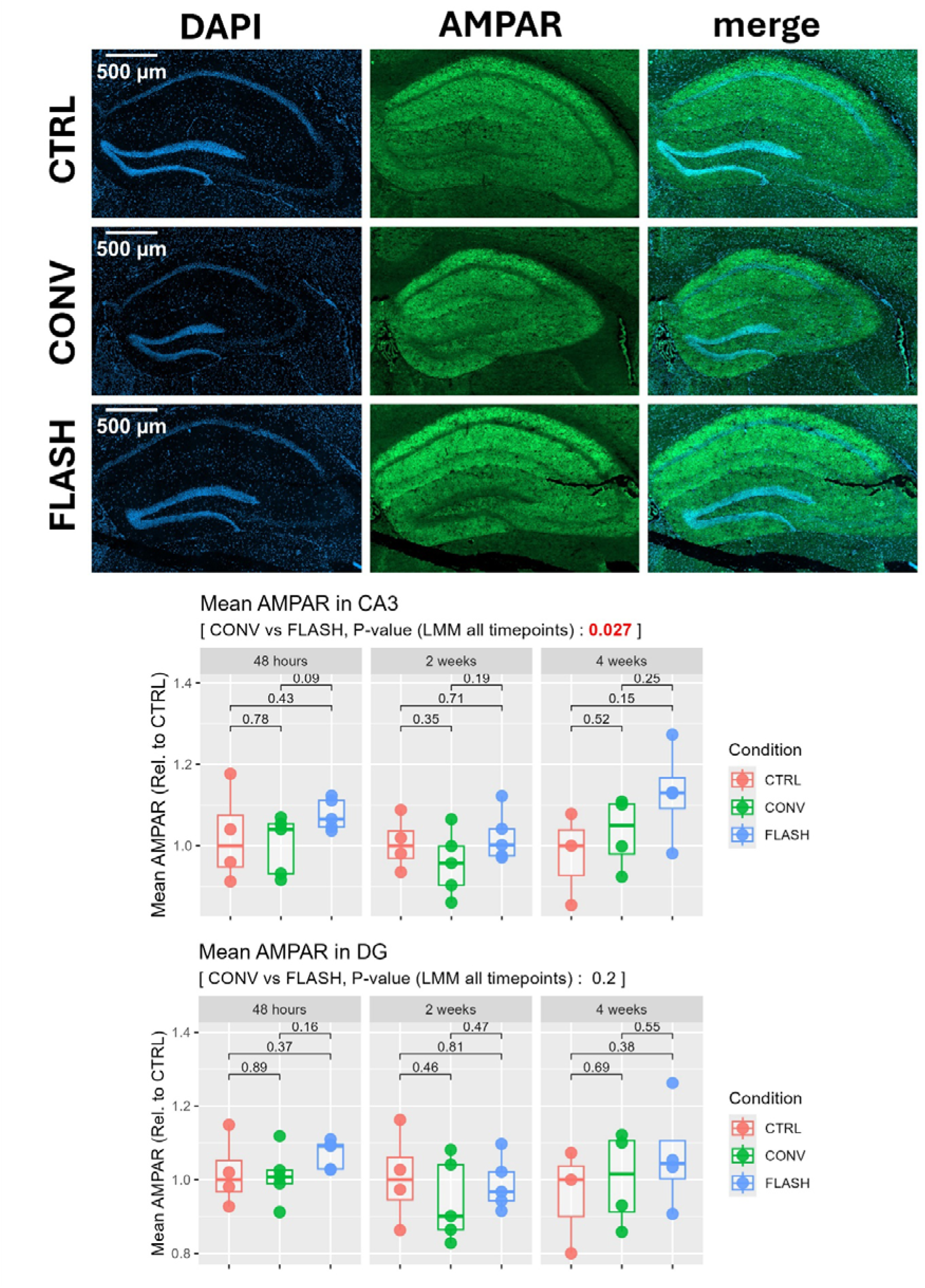
FLASH leads to a sustained AMPAR overexpression 48h, 2 weeks and 4 weeks after irradiation in the CA3. Immunofluorescent staining of hippocampus of control (CTRL), FLASH 10 Gy and CONV 10 Gy irradiated mice. Staining of Gria2 (AMPAR) and nuclei (DAPI). Representative images at the top and quantification across CA3 and DG. Each point represents a different mouse and is the mean of technical replicates (n=3-6). Technical outliers were excluded based on Tukey’s fences method. Pair comparison is obtained from a t-test and assessment over all timepoints was done using a LMM with mouse number and timepoint as random intercepts. See also supp figure 5 and 6.

## Discussion

Our study revealed an early and durable activation of AMPAR following FLASH-RT exposure in hippocampal neurons. This profile involves upregulation of neuronal signals, including synaptic vesicle cycle and LTP pathways as well as key factors known to control synaptic plasticity, with genes encoding for calcium signaling, glutamate receptors and mitochondrial oxidative phosphorylation (OXPHOS). Early transcriptional upregulation of *Gria* translated into increased AMPAR protein levels at 48h in the DG and CA3 region and sustained higher AMPAR expression at 2 and 4 weeks post-FLASH. These findings support the idea that synaptic strength and neurotransmission are preserved over time in FLASH-irradiated samples. We provide three possible and non-mutually exclusive axis of interpretation that might help to explain FLASH-induced neuroprotection: 1) FLASH-RT induces a radiation resistant program in the mature and immature neurons of the CA3, 2) FLASH-RT preserves neural progenitors, 3) FLASH stimulates calcium influx preserving mature/immature neurons and neural progenitors.

Cortical and hippocampal neurons exhibited distinct transcriptional response. In the cortex, the transcriptional program appeared largely dose-rate independent, whereas in the hippocampus it was strongly dose-rate dependent. This suggests <u>region-specific</u> <u>radiosensitivity to dose rate within the brain</u>. This differential sensitivity is consistent with the established relative radio-resistance of the cortex and sensitivity of the hippocampus ^11,12,14^. Moreover, the observation that the FLASH-induced hippocampal transcriptional program resembles the radiation-induced program observed in the cortex, supports the idea that in the CA3 FLASH activates a radioresistant neuronal program. This program involves synaptic function such as LTP, mitochondrial OXPHOS activity and neuronal organization, each of which plays a role in the regulation of synaptic plasticity and is essential for cognitive function. Interestingly, a large proportion of the transcriptional changes in FLASH-irradiated hippocampi occurred after the first fraction of 10 Gy and persisted following subsequent exposure, indicating that the resistance signals could be activated by the first 10 Gy fraction.

While the brain is predominantly composed of terminally differentiated post-mitotic neurons — such as pyramidal cells, GABAergic interneurons, and granule cells — the spatial organization and surrounding microenvironment of these various neuronal populations differ markedly between the cortex and the hippocampus ^15,16^. One possible explanation for the absence of a differential transcriptional signature between FLASH and CONV in cortical neurons could be a transient state of transcriptional inactivity after irradiation. Transcriptional repression is indeed a common stress pattern observed after oxidative stress/hypoxia, energy depletion, excitotoxicity and chronic neuroinflammation all processes induced by irradiation. An alternative hypothesis lies in the cellular composition of the hippocampus, which, unlike the cortex, has a neurogenic niche enriched with undifferentiated, immature progenitor cells. While mature cortical neurons exhibit greater radioresistance, possibly due to their intrinsic resistance to apoptosis ^17^, neural progenitors are highly susceptible to radiation-induced damage at conventional dose rates ^18–20^ but are protected by FLASH ^21,22^. Therefore, the hippocampus-specific transcriptional divergence observed between FLASH and CONV could also reflect the vulnerability of neural progenitors to CONV, which are less sensitive to FLASH and might contribute to the observed transcriptional pattern.

Genes involved in AMPA and NMDA receptor signaling, membrane depolarization, and vesicle recycling (e.g., *Snap25*, *Vamp*, *Syt*, *Dynamin*) were uniquely upregulated following FLASH and AMPAR sustained expression, observed from 48h up to 1-month post-FLASH irradiation, seems to indicate a persistent preservation of synaptic strength and activity. Consistently, we observed an upregulation of *Calm3*, *Ppp3ca*, *Camk2b*, and *Ptk2b* in the hippocampus uniquely after FLASH. These genes are known to play key roles in calcium signaling. They are closely associated with LTP and synaptic vesicle trafficking pathways and our data showed co-expression with AMPAR and NMDAR subunits (*Gria2*, *Grin1*, *Grin2b*). While the resistance molecular cascade is yet to be defined, a working model can be proposed. FLASH may activate calcium signaling via calmodulin (*Calm3*) and calmodulin associated kinase (*Camk2*), leading to Creb phosphorylation and subsequent binding to the Bdnf promoter. This pathway could controls synthesis of new AMPARs and their insertion in the membrane via TrkB activation ^23–26^. Increased AMPAR availability may preserve synaptic plasticity ^27,28^, thereby sustaining long-term potentiation and cognitive function following FLASH irradiation, as previously reported ^4–6^. In parallel, calcium signaling is tightly interconnected with mitochondrial OXPHOS pathways ^29,30^. These findings position calcium signaling as a potential FLASH-specific pathway supporting synaptic function and long-term cognitive preservation and open potentially new avenues of investigation that transcend other organ sites. In this model, an early activation of Camk2 and phosphorylated Camk2 would be expected within the first 24 hours following irradiation. The absence of such findings in our immunofluorescence analyses may reflect the timing of tissue collection, which was performed relatively late (48 hours, 2 and 4 weeks post-irradiation) and may have missed this transient activation window. FLASH modulated calcium influx can also activate several alternative signaling pathways beyond the canonical Camk2–Creb axis. Furthermore, synaptic regulations are complex and sustained activation of excitatory signals might also be neurotoxic, as excessive glutamate activation leads to Ca²□ and neurotransmitter overload ^31^.

In conclusion, FLASH seems to uniquely induce an early and sustained program in the CA3, activating pathways involved in synaptic signaling, mitochondrial function, and the synaptic vesicle cycle. We propose a working hypothesis for a FLASH-induced pathway leading to neuroprotective signals initially activated by calcium influx and signaling, able to induce a radiation-resistance program in neurons and neural progenitors of the CA3.

### Limitations of the study

This study investigated transcriptional profiles in the hippocampi of irradiated mice. The focus on an acute post-irradiation time point (24h) sought to identify initial molecular events triggered by FLASH, but cannot discount additional transcriptional modifications at protracted time points. Although present findings identified early Gria2 induction and sustained AMPAR protein upregulation in the CA3 after FLASH, future experiments are required to dissect the full cascade of events transpiring from early calcium influx and signaling to Bdnf/Creb and mitochondrial OXPHOS. Further work looking at early (10 min-6h post-RT) calcium mediated events can provide for additional assessments of select signaling targets such as pCreb, pErk, pAkt, pCamk2, c-Fos, Arc, Egr1, Bdnf, and mitochondrial Ca²□/OXPHOS. In addition, selective blockade of AMPAR signaling using hippocampal infusion of pharmacological compounds (e.g., NBXQ) will help substantiate the role of AMPAR-mediated neuroprotection. Lastly, a single high dose pulse of FLASH was used to maximize both average and instantaneous dose-rate, a validated method applied across many of our prior FLASH studies of the brain. Notwithstanding, validating present findings with proton and photon FLASH beams with dose rates in the range of 100 Gy/s will support further the clinical translation of FLASH ^32–34^.

## Resource Availability

### Lead Contact

Marie-Catherine Vozenin

### Material availability

Not applicable

### Data and Code Availability

All scripts used for the analysis and plotting can be found at https://gitlab.unige.ch/lirr/nanostring_flash_mouse_brain. Raw and processed data are publicly available in Gene Expression Omnibus (GEO) under accession number GSE305149. Other data can be made available upon request.

## Other

The authors thank Dr V. Grilj for irradiation procedures and dosimetry, Epalinges animal facility, Dr S. Tissot team at the immune landscape laboratory Platform UNIL/CHUV, Bioimaging core facility UNIGE, Dr J. Prados from bioinformatics support platform UNIGE, P. Angelino from Translational data science SIB, Lausanne. Graphical abstract was done with Biorender.

## Credit authorship contribution statement

**Vozenin Marie-Catherine** Conceptualization – Investigation – Supervision – Writing original draft – Securing fundings. **Louis Kunz** Investigation (bio-informatic analysis)–Methodology – Writing original draft. **Aymeric Almeida** Investigation (animals and DSP)–Methodology – Writing original draft. **Michèle Knol** Investigation (IHC). **Benoit Petit** Investigation (irradiation and animal follow up) – Methodology. **Eniko A. Kramár, Marcelo A. Wood** writing original draft. **Charles L. Limoli** Conceptualization – Supervision – Writing original draft. **All authors** Write, Review & Edit.

## STAR Methods

### Experimental Model

All animal procedures were conducted in accordance with the Swiss ethics committee (VD3603) for animal experimentation. Eighteen eight-week-old female C57Bl/6J mice were purchased from Charles River Laboratories and were irradiated at 10 weeks of age.

### Whole Brain Irradiations

Irradiation was performed using a prototype 6 MeV electron beam, Oriatron type 6e (eRT6; PMB Alcen), LINAC, available at Lausanne University Hospital and described previously ^35^. Dosimetry has been extensively described and published to ensure reproducible and reliable biological studies ^36^. Animals were randomized in 5 groups and were sham-, for controls, or whole brain irradiated using a single dose of 10 Gy or 3 fractions of 10 Gy delivered 48 hours apart. A 17-mm graphite applicator was used and CONV was delivered at 0.09 Gy/s whereas FLASH was delivered in 1 pulse of 5.6×10^6^Gy/s, maximizing both instantaneous and average dose-rate. 24h post-treatment completion, whole brains were collected and stored at 4°C in 4% PFA for 48 hours before Formalin-Fixed Paraffin-Embedded (FFPE) before processing.

For follow-up immunohistochemistry, C57Bl/6J female twelve-week-old mice were irradiated using the same protocol and a single dose of 10 Gy before cryopreservation of sampled brains, 48 hours, 2 weeks and 4 weeks post irradiation.

### GeoMx DSP RNA profiling in situ hybridization

FFPE tissue section of 5-µm were processed according to the provider standard protocol for the GeoMx Digital Spatial Profiler (DSP) ^37^. Neurons were identified using NeuN (1:100, Millipore ABN78) and revealed with a secondary goat anti-rabbit Alexa 594 (1:1000, Abcam ab150080). Vessels and astrocytes were also stained using an antibodies cocktail composed of CD31 Alexa 488 (Vessels, 1/100, R&D FAB3628G) and GFAP Alexa 647 (Astrocytes, 1/100, Novus NBP2-3318). The nuclear marker Syto83 (Invitrogen 511364) was also used. Whole slides were imaged at x20 magnification and morphologic markers were used to select regions of interest (ROI). For each mouse 2 ROIs were selected in the Cortex and 4 in the hippocampus, specifically 1 in the CA1, 1 in the CA3, 1 in the DG and 1 in the SO. Automatic segmentation of ROIs based on CD31, NeuN and GFAP markers were performed to define area of illumination (AOIs). AOIs were exposed to 385□nm light (UV), releasing the indexing oligonucleotides to be sequenced. Sequencing libraries were generated by PCR from the photo-released indexing oligos. Pooled libraries were single-sequenced at 27 base pairs and with the single-index workflow on an Illumina NovaSeq S4 instrument.

### Immunohistochemistry

Brains of 10 Gy irradiated mice were coronally sectioned to 10µm around the hippocampal region. Slides were fixed in PFA and Tp citrate was used for antigen retrieval. Goat F(ab) Anti-Mouse (Abcam ab6668) was applied on Camk2 and pCamk2 staining, and PBS+DKS 4%+triton 0.4% were used as blocking solution on all tissues. Primary antibodies were diluted 1:200 in PBS+DKS 1%+triton 0.4 % and applied overnight (Rabbit anti-GluA2/Gria2 Alomone AGC-005; Guinea pig anti-GluN1/Grin1 Alomone A21435; Rabbit anti-Camk2 alpha Abcam AB131468; Mouse anti-Phospho-CaMKII alpha (Thr286) Invitrogen MA1-047). Secondary antibodies were diluted 1:500 in the same solution and applied for 60min (AF555 Dk-aMs Life Technologies A31570, AF488 Dk-aRb Life Technologies A21206, AF555 Gt-aGp Invitrogen A21435). Mounting was done using Gold antifade reagent with DAPI (ProLong^TM^ Invitrogen P36935). Slide acquisition was done using Zeiss Axioscan.Z1 at the bioimaging facility of UNIGE and image analysis was done in QuPath ^38^ before downstream analysis in R ^39^ using ggplot2 ^40^. Technical outliers were excluded based on Tukey’s fences method. Pair comparison is obtained from a t-test and assessment over all time points was done using a LMM with mouse number and timepoint as random intercepts.

### Bioinformatic analysis

DSP data were analyzed in R 4.4.0 (2024) ^39^ according to Nanostrings recommendations ^41^, using their in-house tools ^42–44^, with the exception of normalization, which was performed using the trimmed mean of M values (TMM) algorithm of edgeR ^45,46^. Other packages were used for data manipulation and plotting ^40,47–51^ as well as gene set enrichment analysis ^52–55^.

First, quality control (QC) was performed for the segments using a set of thresholds on chosen descriptive statistics of the segment (Table supp 1). Chosen parameters were used as recommended by Nanostring. Then, a Grubb test was performed for gene QC, also using parameters recommended by Nanostring (Table supp 1). Next, negative probes were used to evaluate the background of quantification and define limit of quantification thresholds for filtering. Segments that contain less than 5% of the genes above the gene detection threshold and genes that are found in less than 5% of the segments were removed from the analysis. After filtering, normalization was performed using the TMM method of the edgeR package. Robust gene detection rate within segments and cell type specificity were confirmed, enabling downstream analysis (Figure supp 1A-C)

For unsupervised analysis, uniform manifold approximation and projection (UMAP) ^56^ and t-distributed stochastic neighbor embedding (tSNE) ^57^ were used to perform dimension reduction.

For figure 1, 2 and supp figure 2, where the brain regions contain more than one ROI (i.e. 2 for cortex, 4 for hippocampus) data were not aggregated, each ROI was analyzed independently. For all other figures, CA1, CA3, DG, SO, only one ROI was segmented per region per mouse, thus each ROI is fully independent. Differential gene expression analysis was performed using a linear mixed model (LMM) with the irradiation modality set as the test variable and the slide number as the random intercept. To prevent batch effects, one CTRL, one CONV and one FLASH brain section was loaded on each slide to be analyzed.

The metric -sign(FC) • log2(pvalue) was used to rank genes, allowing gene set enrichment analysis (GSEA) to be performed using the disease ontology semantic and enrichment (DOSE) ^53^ analysis, implemented in clusterprofiler ^52^. For each pathway a normalized enrichment score (NES) and a p-value were obtained.

All scripts used for the analysis and plotting can be found at https://gitlab.unige.ch/lirr/nanostring_flash_mouse_brain. Raw and processed data can be found at NIH GEO under accession number GSE305149. All key resources can also be found in a summary table according to STAR methods.

---

**Fig. supp 1:**
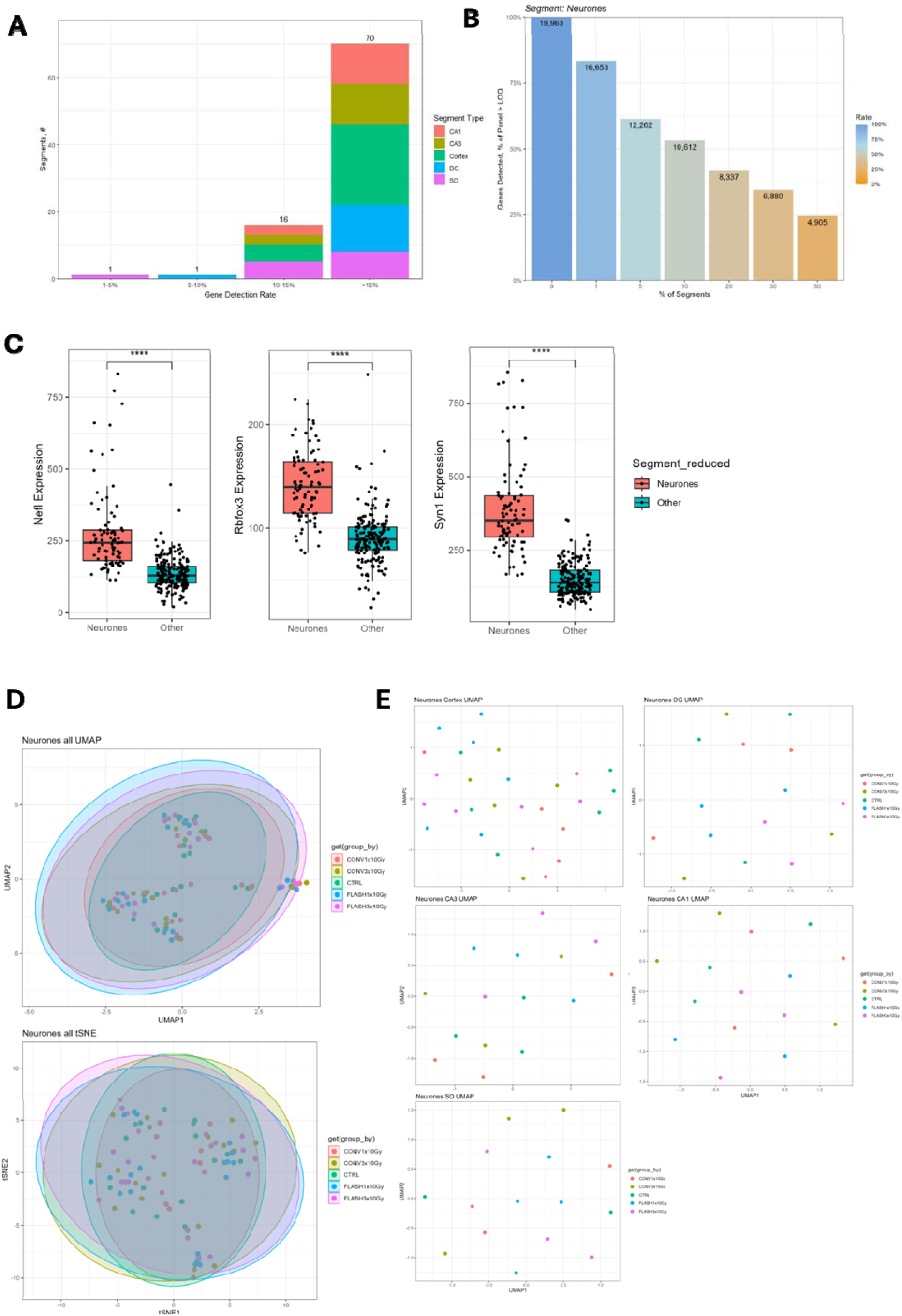
Confirmation of robust gene detection rate within segments and confirmation of cell type specificity. (A) Histogram of number of ROI (segments) for increasing gene detection rate thresholds colored per subregions. (B) Percentage of genes detected i.e. above limit of quantification (LOQ). (C) Expression of specific markers of neurons across different cell types (as identified with immunostainings). P-values computed using a Mann-Whitney test. One, two, three and four stars respectively representing p-values smaller than 0.05, 0.01, 0.001, 0.0001. (D) UMAP and tSNE colored per irradiation conditions for all regions (all) or (E) individual regions considered (Cortex, DG, CA3, CA1, SO). Related to STAR methods and Figure 1.

**Fig. supp 2:**
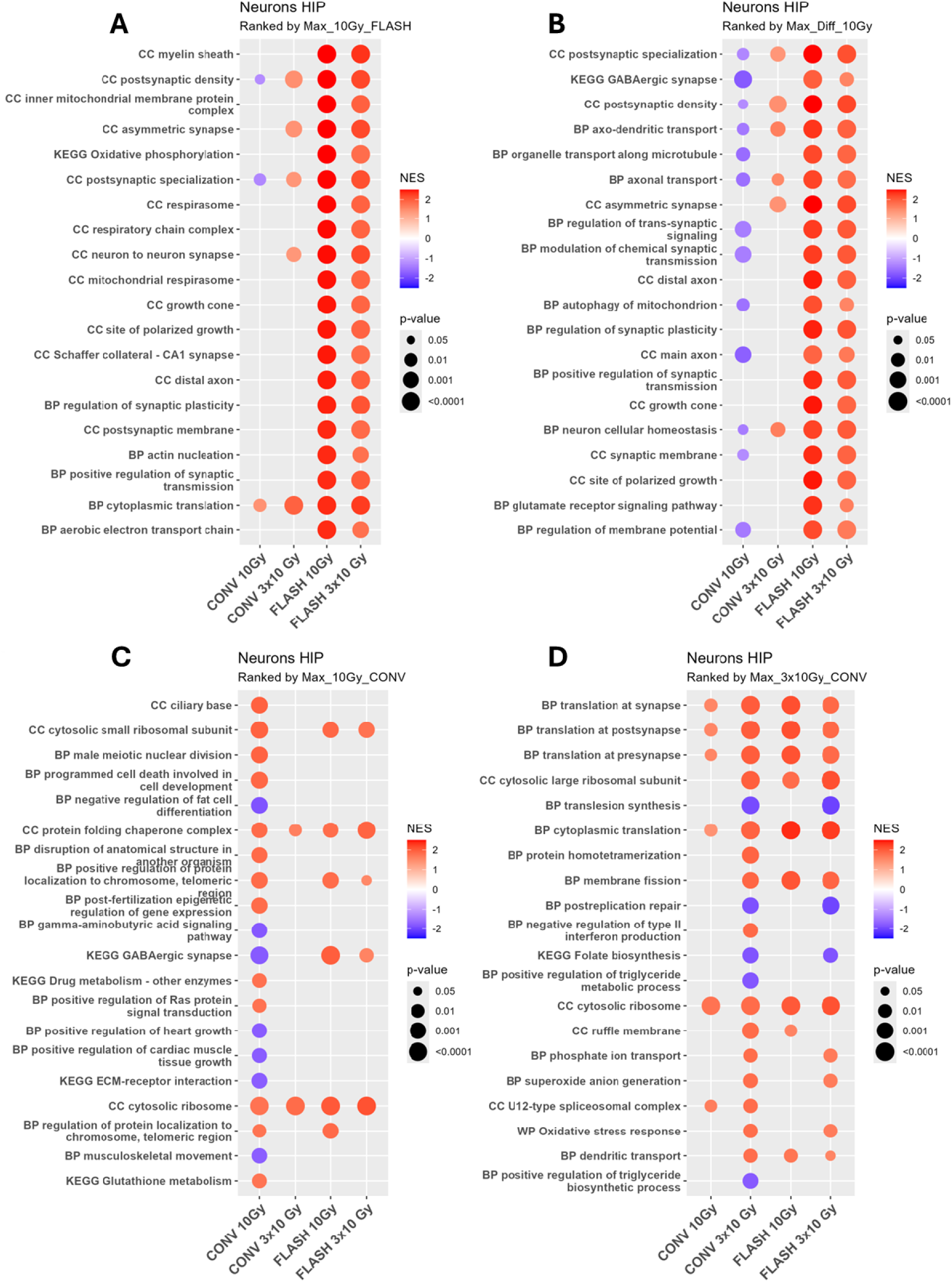
CONV presents dose dependent imprint in the hippocampus while FLASH seems to induce a dose agnostic response. (A) Output of GSEA on Hippocampus signature, pathways are ranked according to the maximum absolute NES of FLASH 10 Gy (B) the maximum absolute difference in NES between CONV 10 Gy and FLASH 10 Gy, (C) maximum absolute NES of CONV 10 Gy (D) or maximum absolute NES of CONV 3×10 Gy. Normalized enrichment score (NES) as the color and p-value as the dot-size.. Related to Figure 2.

**Fig. supp 3:**
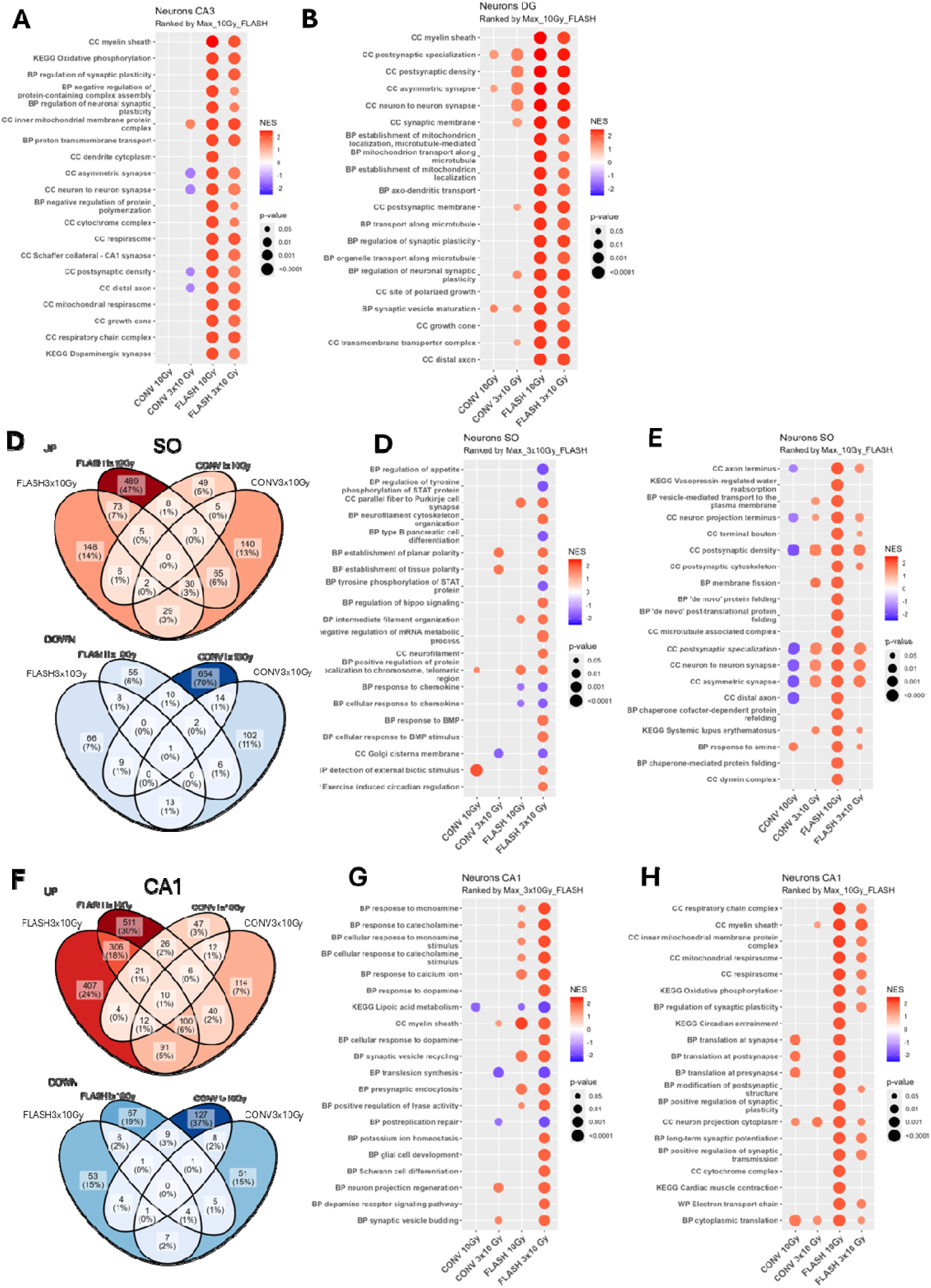
While in the CA3 and the DG FLASH induces a dose agnostic response, in the SO and in the CA1 the imprint always depends on the dose. (A) Output of GSEA signature, pathways are ranked according to the maximum absolute NES of FLASH 10 Gy for the CA3 (B) or the DG. (C) Venn Diagrams of Up (NES > 0, p-value < 0.05) and Down (NES < 0, p-value < 0.05) regulated pathways in the SO. (D) Output of GSEA on SO signature, pathways are ranked according to the maximum absolute NES of FLASH 3×10 Gy (E) or max absolute NES of FLASH 10 Gy. (F) Venn Diagrams of Up (NES > 0, p-value < 0.05) and Down (NES < 0, p-value < 0.05) regulated pathways in the CA1. (G) Output of GSEA on CA1 signature, pathways are ranked according to the maximum absolute NES of FLASH 3×10 Gy (H) or max absolute NES of FLASH 10 Gy. Related to Figure 3.

**Fig. supp 4:**
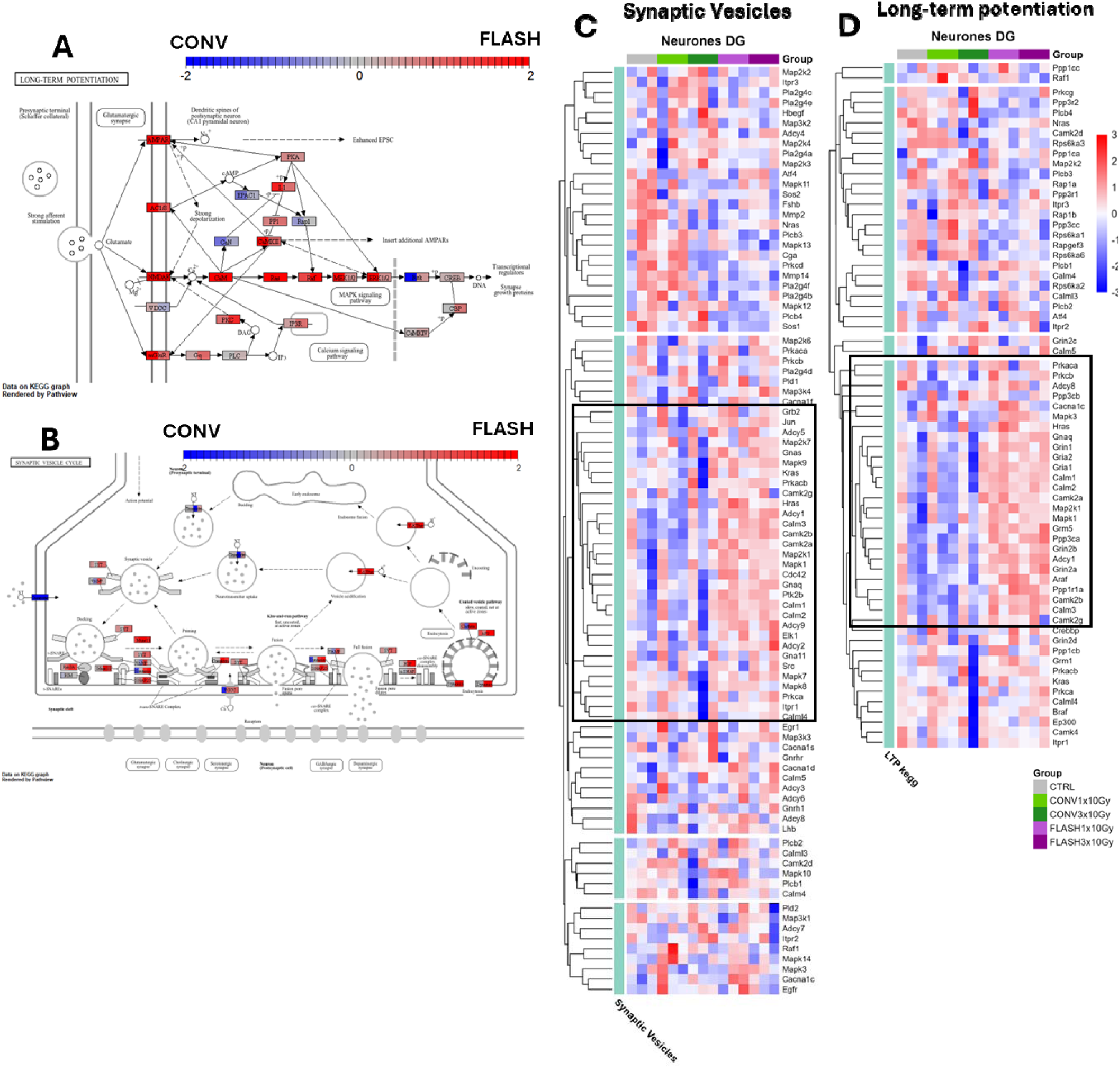
Pathway elements and specific genes show a similar pattern in the DG as in the CA3 for long term potentiation and synaptic vesicle cycle. (A) Pathways diagrams made using *Pathview* for KEGG long-term potentiation (B) and synaptic vesicle cycle based on DG imprint. Color is - an arbitrary scale of enrichment from the highest expression in CONV (-2) to the highest expression in FLASH (+2). Each element of the pathway is split across center for CONV/FLASH comparison at 10 Gy (left) and at 3×10 Gy (right). (C) Gene regulation heatmap showing for neurons of DG, with black squares drawn arbitrarily for KEGG synaptic vesicles (D) and long-term potentiation.Related to Figure 4 and 5.

**Fig. supp 5:**
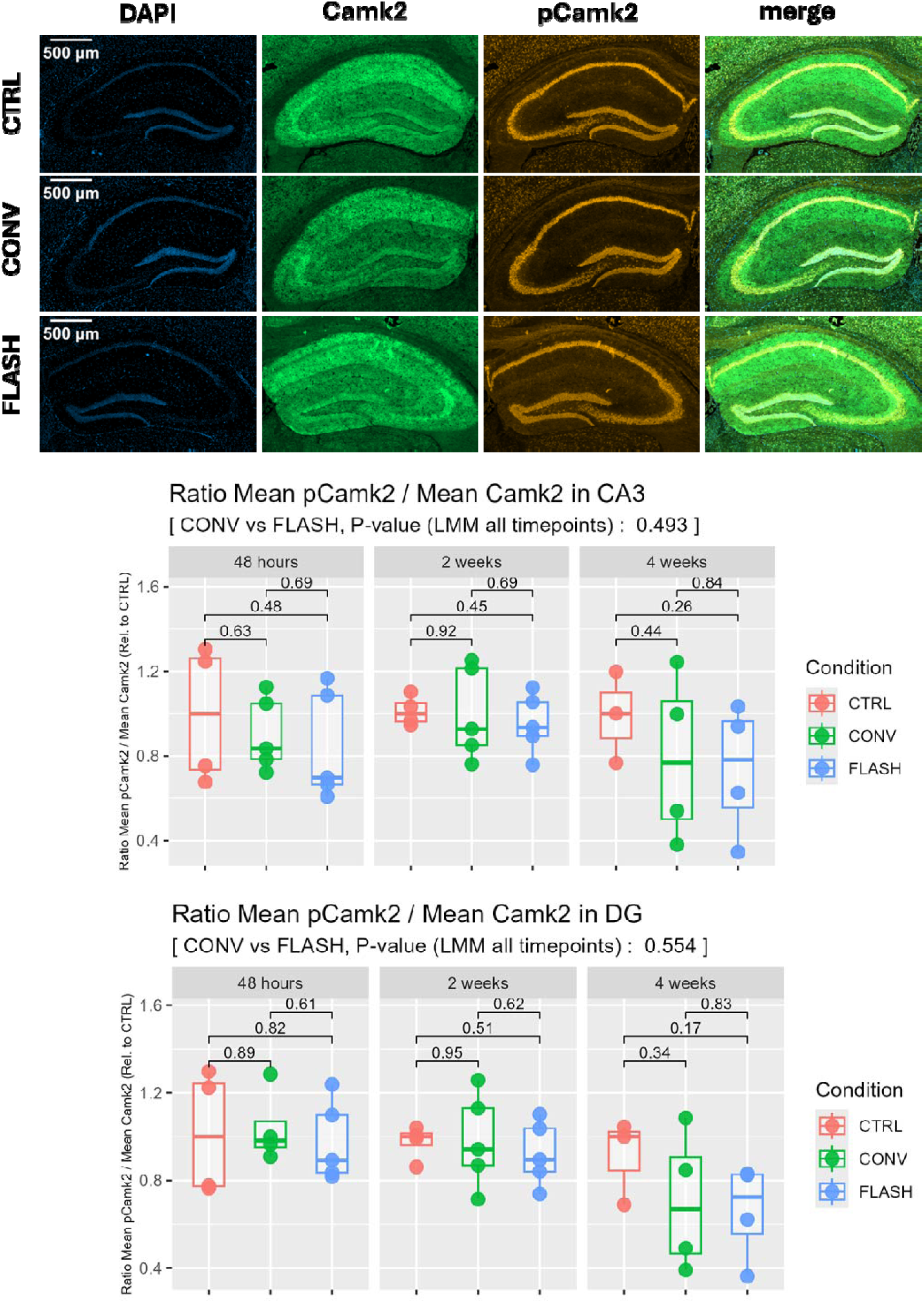
Phosphorylated Camk2 ratio to Camk2 shows steady levels 48h, 2 weeks and 4 weeks after irradiation. Immunofluorescent staining of hippocampus of control (CTRL), FLASH 10 Gy and CONV 10 Gy irradiated mice. Staining of phosphorylated *Camk2* (p*Camk2*), *Camk2* and nuclei (DAPI). Representative images at the top and quantification across DG or CA3 at the bottom. Each point represents a different mouse and is the mean of technical replicates (n=3-6). Technical outliers were excluded based on Tukey’s fences method. Pair comparison is obtained from a t-test and assessment over all timepoints was done using a LMM with mouse number and timepoint as random intercepts.. Related to Figure 5.

**Fig. supp 6:**
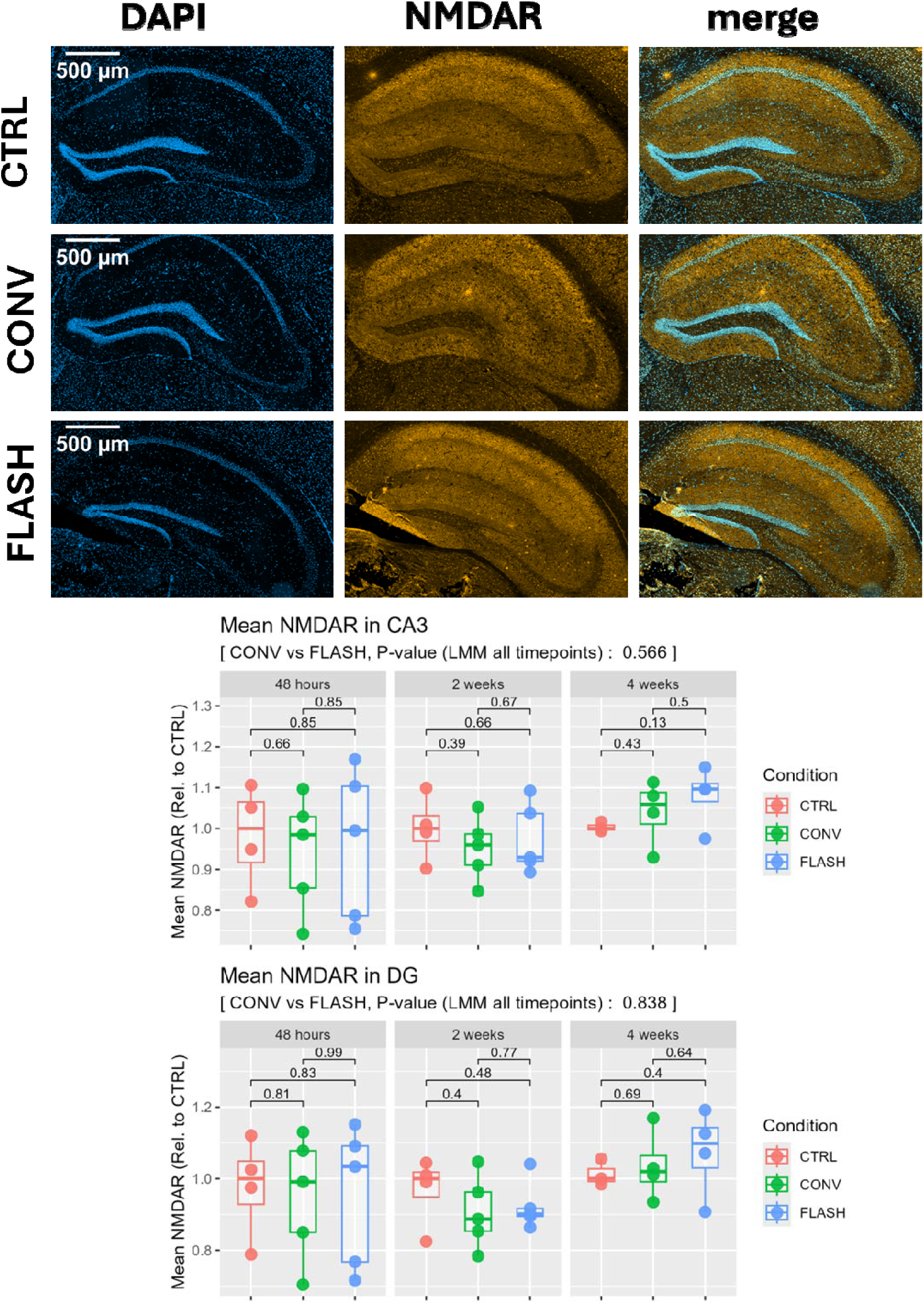
NMDAR are maintained at constant level 48h, 2 weeks and 4 weeks after irradiation. Immunofluorescent staining of hippocampus of control (CTRL), FLASH 10 Gy and CONV 10 Gy irradiated mice. Staining of *Grin1* (NMDAR) and nuclei (DAPI). Representative images at the top and quantification across DG or CA3 at the bottom. Each point represents a different mouse and is the mean of technical replicates (n=3-6). Technical outliers were excluded based on Tukey’s fences method. Pair comparison is obtained from a t-test and assessment over all timepoints was done using a LMM with mouse number and timepoint as random intercepts.Related to Figure 6.

**Supplementary table 1:** DSP Nanostring GeoMX quality check parameters used.

| QC segments |  |
| --- | --- |
| Minimum number of reads | 1000 |
| Minimum % of reads trimmed | 80 |
| Minimum % of reads stitched | 80 |
| Minimum % of reads aligned | 75 |
| Minimum sequencing saturation | 50 |
| Minimum negative control counts | 1 |
| Minimum # of nuclei estimated | 20 |
| Minimum segment area | 1000 |
| QC genes Grubb test |  |
| minimum Probe Ratio | 0.1 |
| percent Fail Grubbs | 20 |

